# Loss of Polycomb Repressive Complex 2 function causes asparaginase resistance in T-acute lymphoblastic leukemia through decreased WNT pathway activity

**DOI:** 10.1101/2023.08.04.552014

**Authors:** Thomas Lefeivre, Cosmin Tudose, Theodora-Ioana Grosu, Luke Jones, Theresa E. León, Kieran Wynne, Giorgio Oliviero, Owen P. Smith, Amélie Trinquand, Marc R. Mansour, Colm J. Ryan, Jonathan Bond

**Author notes:** **Corresponding author:** Prof. Jonathan Bond, Systems Biology Ireland, School of Medicine, University College Dublin, Belfield, Dublin 4, Ireland.

## Abstract

Loss-of-function mutations and deletions in core components of the epigenetic Polycomb Repressive Complex 2 (PRC2) are associated with poor prognosis and treatment resistance in T-acute lymphoblastic leukemia (T-ALL). We leveraged clinical mutational and transcriptional data to identify a functional link between PRC2 alterations and changes in WNT signaling pathway activity in leukemia cells. Computational integration of transcriptomic, proteomic and phosphoproteomic data from an isogenic T-ALL cellular model revealed reduced activity of the WNT-dependent stabilization of proteins (WNT/STOP) pathway in cells lacking core PRC2 factor EZH2. We discovered that PRC2 loss significantly reduced sensitivity to key T-ALL treatment asparaginase, and that this was mechanistically linked to increased cellular ubiquitination levels that bolstered leukemia cell asparagine reserves. We further found that asparaginase resistance in PRC2-depleted leukemic blasts could be mitigated by pharmaceutical proteasome inhibition, thereby providing a novel and clinically tractable means to tackle induction treatment failure in high-risk T-ALL.

## Introduction

Outcomes for young patients with T-acute lymphoblastic leukemia (T-ALL) have lagged behind cases of B-ALL in the same age group, in part due to a less complete mechanistic understanding of the molecular drivers of treatment resistance.

Recent large scale genomic efforts have highlighted a key role for epigenetic factors in aggressive T-ALL biology, as shown by the high frequency of loss-of-function mutations and deletions in genes coding for Polycomb Repressive Complex 2 (PRC2) factors in the immature T-ALL subgroup (1,2). In line with the elevated rates of remission induction failure in these cases (3), reduced PRC2 function has been shown to correlate with chemotherapy resistance in both T-ALL and acute myeloid leukemias (AML) that harbor a similar mutational genotype in core PRC2 components *EZH2*, *EED* and *SUZ12* (1,4,5). Various molecular explanations have been proposed to explain chemoresistance in PRC2-deficient T-ALL, including changes in NOTCH1-mediated transcriptional regulation (6) and increased mitochondrial apoptosis resistance (7), but a clear mechanistic link between PRC2 loss of function and altered sensitivity to any specific T-ALL remission induction treatment has not yet been established.

Clues to the mechanistic basis of T-ALL therapy resistance may be revealed through careful analysis of mutational co-occurrence, which can inform a better understanding of interplay between apparently discrete molecular pathways. Epigenetic alterations in T-ALL frequently segregate with activating mutations in signaling molecules, suggesting that these alterations may interact functionally (2). Signaling pathways are appealing treatment targets due to their therapeutic tractability, and there is evidence that kinase inhibition can be leveraged to improve drug responses in combination with epigenetic agents (8), and in leukemias that harbor epigenetic factor alterations (9).

Recent work has revealed important roles for signaling pathways that are comparatively less studied in T-ALL biology. In particular, several reports have elegantly identified that therapeutic responses to the key T-ALL remission induction treatment asparaginase are heavily influenced by the activity level of the WNT-dependent stabilization of proteins (WNT/STOP) pathway, which regulates leukemic blast asparagine reserves (10,11).

In this study, we leveraged clinical mutational and transcriptional data and cellular models of leukemia PRC2 loss to explore the functional interplay between epigenetic factor and signaling pathway alterations in T-ALL, with an aim of understanding how these interactions alter responses to specific therapies.

## Results

### Mutual exclusivity and co-occurrence between epigenetic factor and signaling pathway genetic alterations in T-ALL

Patterns of co-occurrence and mutual exclusivity between genetic alterations can reveal biological functional relationships in cancer cells. For example, somatic mutational co-occurrence may reveal synergies that promote oncogenesis, while mutual exclusivity can identify gene pairs that are synthetic lethal, such that their combined alteration causes a significant fitness defect. Identification of these genetic interactions may therefore inform treatment approaches by revealing tumor-specific molecular vulnerabilities (12).

To evaluate such relationships in T-ALL, we analyzed copy number profile and somatic mutation data from a published cohort of 249 pediatric T-ALL cases (2). We restricted our analysis to alterations of genes that encode either signaling proteins or epigenetic regulators, and used an established approach (13) to evaluate all possible co-occurrence and mutual exclusivity relationships within and between these gene sets. All statistically significant (p value < 0.05, FDR < 0.25) results of these analyses are shown in Supplemental Table S1. These results included identification of previously reported relationships, such as mutual exclusivity between mutations of *PTEN* and *WT1*, and co-occurrence between *JAK1* and *JAK3* mutations (14,15).

In contrast with prior studies that focused on genetic interactions between pairs of mutations, the fact that our analyses included both mutation and copy number data (full details in Supplemental Figure S1A) allowed us to detect a number of previously unreported interactions between genes encoding signaling molecules and epigenetic proteins (Supplemental Figure S1B). In particular, we identified several associations involving PRC2 components and factors related to the Wingless/Integrated (WNT) signaling pathway (Figures 1A and 1B) that led us to explore these interactions in more depth in later experiments. For instance, *EZH2* mutation co-occurred with the shallow loss of a group of six genes located on 17q11.2, including *NLK*, which encodes Nemo-like kinase, a negative regulator of WNT signaling (16). *EZH2* mutation also frequently co-occurred with deep loss of *CDKN2A* that modulates WNT signaling as part of its tumor suppressor activity (17). We further found that mutation of *EZH2* was mutually exclusive with mutation of *FBXW7*, which alongside its role in NOTCH pathway signaling, antagonizes WNT activity through β-catenin degradation (18). Taken together, these genetic interaction patterns raise the possibility of a hitherto undetected functional link between WNT signaling and epigenetic factors in T-ALL, and between WNT and PRC2 core activity specifically.

**Figure 1:**
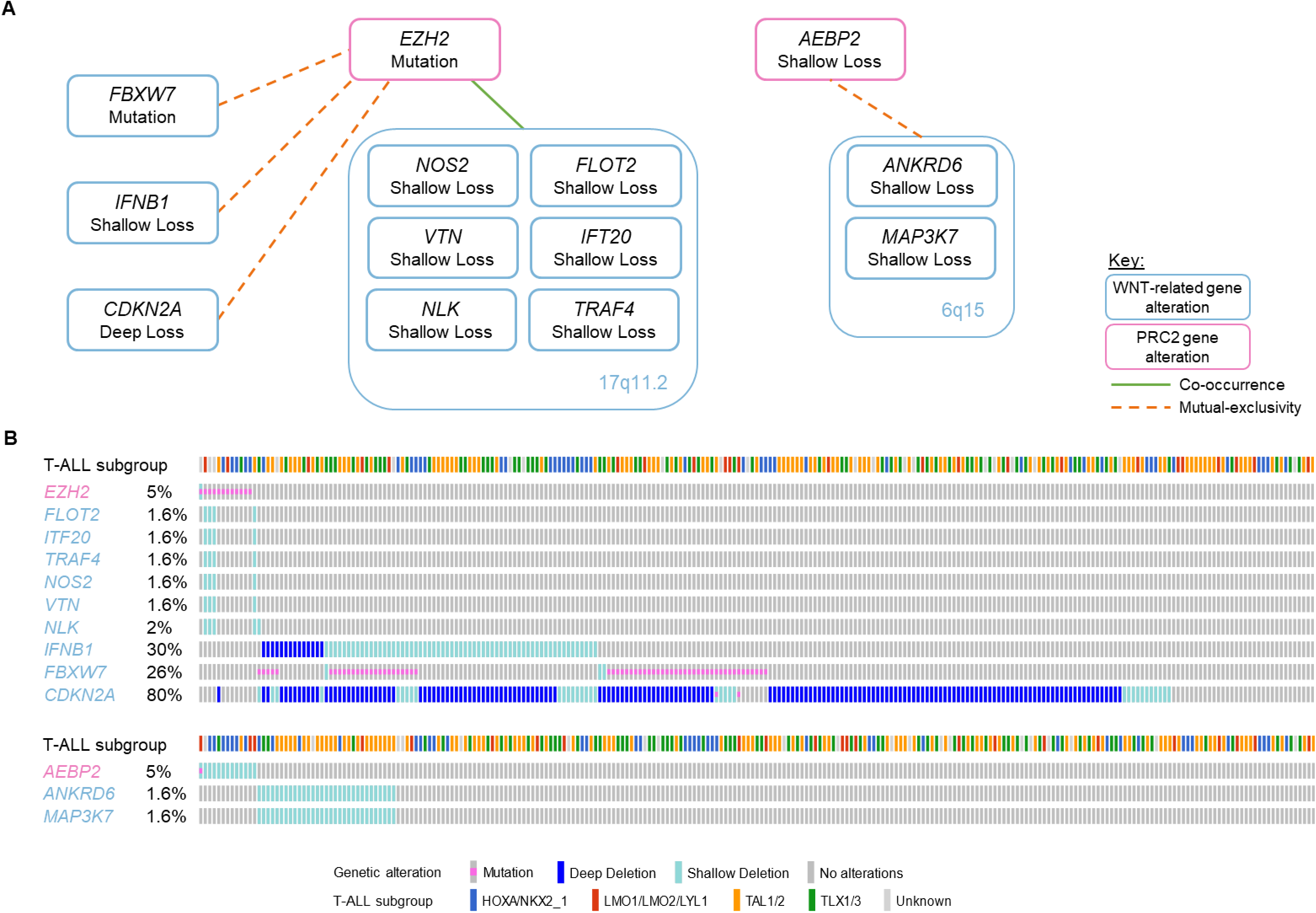
Mutual exclusivity and co-occurrence between epigenetic factor and WNT pathway genetic alterations in T-ALL. **(A)** Graphical representation of statistically significant interactions between genes coding for PRC2 factors (pink boxes) and WNT signaling components (blue boxes), as determined by DISCOVER analysis (13) of a T-ALL cohort. **(B)** Oncoprint of significant associations showing co-occurrence and mutual exclusivity of mutations, deep deletions, and shallow deletions in individual cases. T-ALL subtype corresponds with annotation from the original description of this cohort (2).

### Loss of PRC2 function correlates with reduced transcription of WNT pathway targets

Given the multiple associations between epigenetic and signaling factor genetic alterations identified in our analysis, we next set out to characterize the molecular effects of PRC2 loss on signaling pathways in T-ALL. To do so, we made an isogenic Jurkat cell line model using CRISPR/Cas9 to target exon 3 of *EZH2* (Figure 2A, Supplemental Figure S2A).

**Figure 2:**
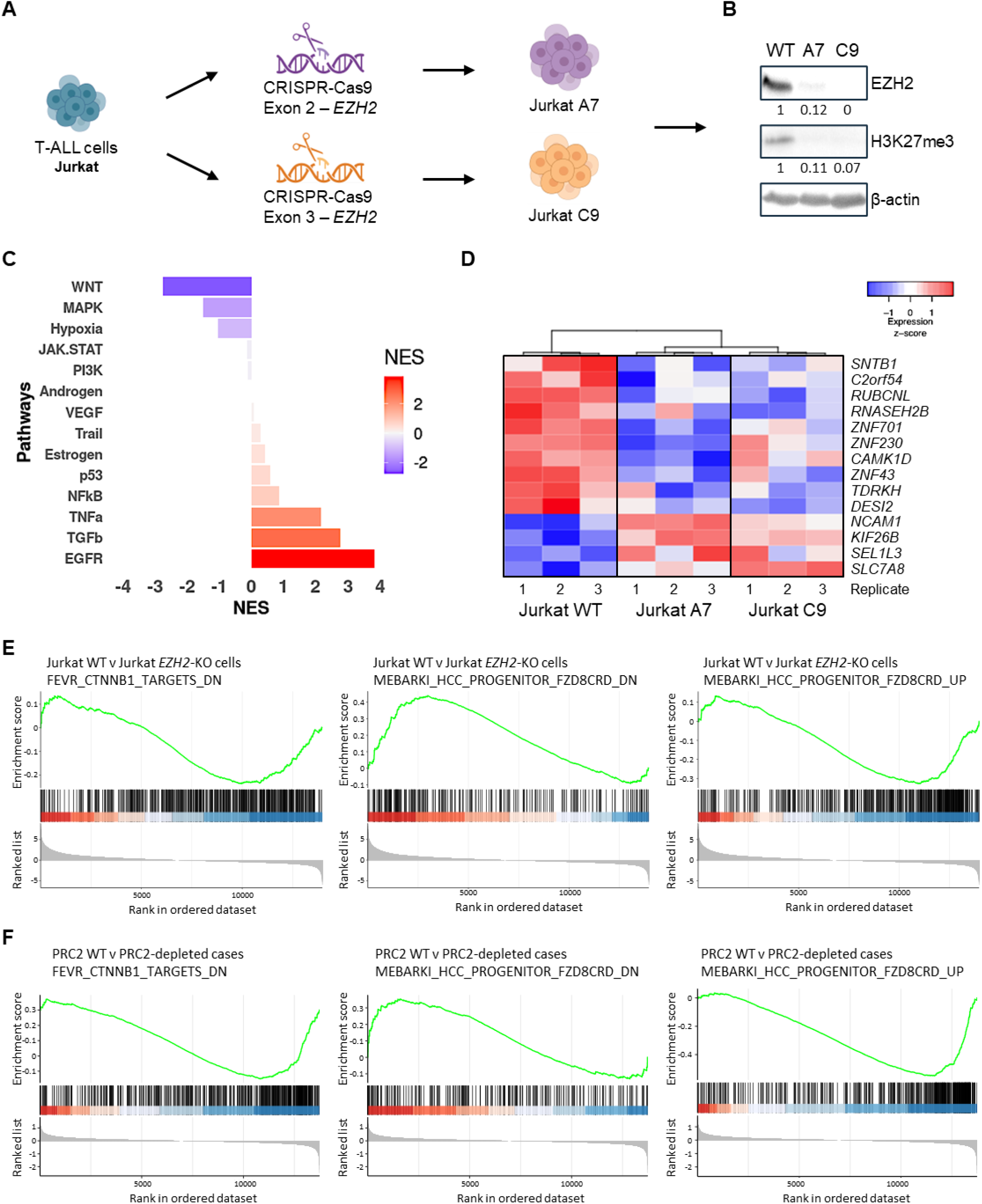
Loss of PRC2 function correlates with reduced transcription of WNT pathway targets. **(A)** Schema of isogenic cell line model and **(B)** confirmation of reduced EZH2 and H3K27 trimethylation (me3) in EZH2 knockout (KO) clones A7 and C9 by protein immunoblotting. **(C)** PROGENy analysis of signaling pathway activity from RNA-seq transcriptional readouts. Enrichment in EZH2-KO cells is depicted, i.e., negative enrichment corresponds to predicted decreased activity of that pathway in PRC2-depleted cells. NES: Normalized enrichment score. See Figure S2D for comparison of individual clones with WT samples. **(D)** Altered transcription of WNT pathway targets in PRC2-depleted T-ALL. Genes with statistically significant gene expression changes between WT and EZH2-KO Jurkat cell lines are shown. Gene set enrichment analyses (GSEA) of WNT pathway-related signatures in **(E)** EZH2 WT and KO Jurkat cells and **(F)** PRC2 WT and altered primary T-ALLs reveal similar transcriptional patterns. Gene sets are indicated above each panel and correspond to β-catenin (left panels) and Frizzled (middle and right panels) targets. See also Supplemental Figure S2F.

As CRISPR/Cas9 knockout (KO) clones are known to harbor cryptic genetic and functional heterogeneity (19), we included an additional *EZH2* KO clone that was previously generated from a different parental Jurkat line using an alternative gene editing strategy (20) in our analyses, to increase the likelihood that any molecular changes we detected were specifically related to PRC2 loss, rather than random clonal phenotypic variability. We confirmed reductions of EZH2 protein expression and function by measurement of trimethylation of lysine 27 of histone H3 (H3K27me3) in these clones by immunoblotting (Figure 2B) before performing comprehensive molecular characterization of both lines.

We reasoned that transcriptional readouts would provide the best starting point for assessing changes in cellular signaling activity following EZH2 loss, and so prioritized these analyses. We performed RNA-sequencing of wild-type (WT) and EZH2-deficient lines and used PROGENy (Pathway RespOnsive GENes) (21) to analyze the results. PROGENy uses curated gene expression and perturbation data to infer the activity of 14 separate signaling cascades, thereby helping us to assess how the function of these pathways differed between WT and EZH2-depleted cells. As the two EZH2-KO clones presented similar gene expression profiles (Supplemental Figure S2B), these transcriptional results were combined and compared with WT readouts for the analysis.

PROGENy revealed altered activity of multiple signaling pathways in EZH2-null cells (Figure 2C and Supplementary Figure S2C), including EGFR, TGFβ, TNFα (all increased), MAPK, and hypoxia (both decreased). Given the genetic interactions between *EZH2* and WNT factors that we discovered in T-ALL cases (Figure 1), we were particularly struck by the differences in inferred WNT pathway activity between WT and EZH2-deficient cells. WNT was found to be the most downregulated signaling cascade in EZH2-KO cells, with marked transcriptional changes in targets of both canonical and non-canonical WNT pathways (Figure 2D and Supplemental Figure S2D).

To confirm that these gene expression changes were not specific to the Jurkat cell line model, we analyzed the same T-ALL cohort (1) as was used for the DISCOVER analysis, using gene set enrichment analysis (GSEA) to determine the transcriptional differences between PRC2-WT and PRC2-altered cases (cohort details are in Supplemental Figure S2E). Initial review of the most significantly altered oncogenic signatures (C6) gene sets from MSigDB confirmed marked negative enrichment for genes previously described to decrease in expression following either EZH2 or EED knockdown in samples with PRC2 loss, providing support for use of this method to determine transcriptional differences between WT and PRC2-altered primary samples (Supplemental Figure S2F).

Strikingly, several sets of genes related to WNT pathway activity were also among the most enriched gene sets in PRC2-altered cases in this unbiased analysis, including WNT3A/Frizzled, β-catenin, and LEF1 targets. In each case, transcriptional differences between PRC2-altered and PRC2-WT patient T-ALL samples corresponded with those seen between WT and EZH2-null Jurkat cells (Figures 2E and 2F and Supplemental Figure S2F). Taken together, these results provide transcriptional evidence of decreased WNT activity in primary leukemias with altered PRC2 function, and suggest that the molecular changes in the Jurkat EZH2-KO model can be extrapolated to the PRC2-depleted T-ALL molecular landscape *in vivo*.

### Multiomics analysis confirms reduced WNT signaling activity in PRC2-depleted T-ALL

We next determined whether the transcriptional changes associated with PRC2 loss correlated with proteomic and phosphoproteomic assessment of signaling pathway activity.

Mass spectrometric analysis of whole cell extracts revealed significant differences in the proteomes of WT and EZH2-deficient cells (Figure 3A). Importantly, there was marked overlap in the repertoire of proteins that were significantly up-regulated and down-regulated in each EZH2-KO clone, compared with the WT sample (Supplemental Figure S3A).

**Figure 3:**
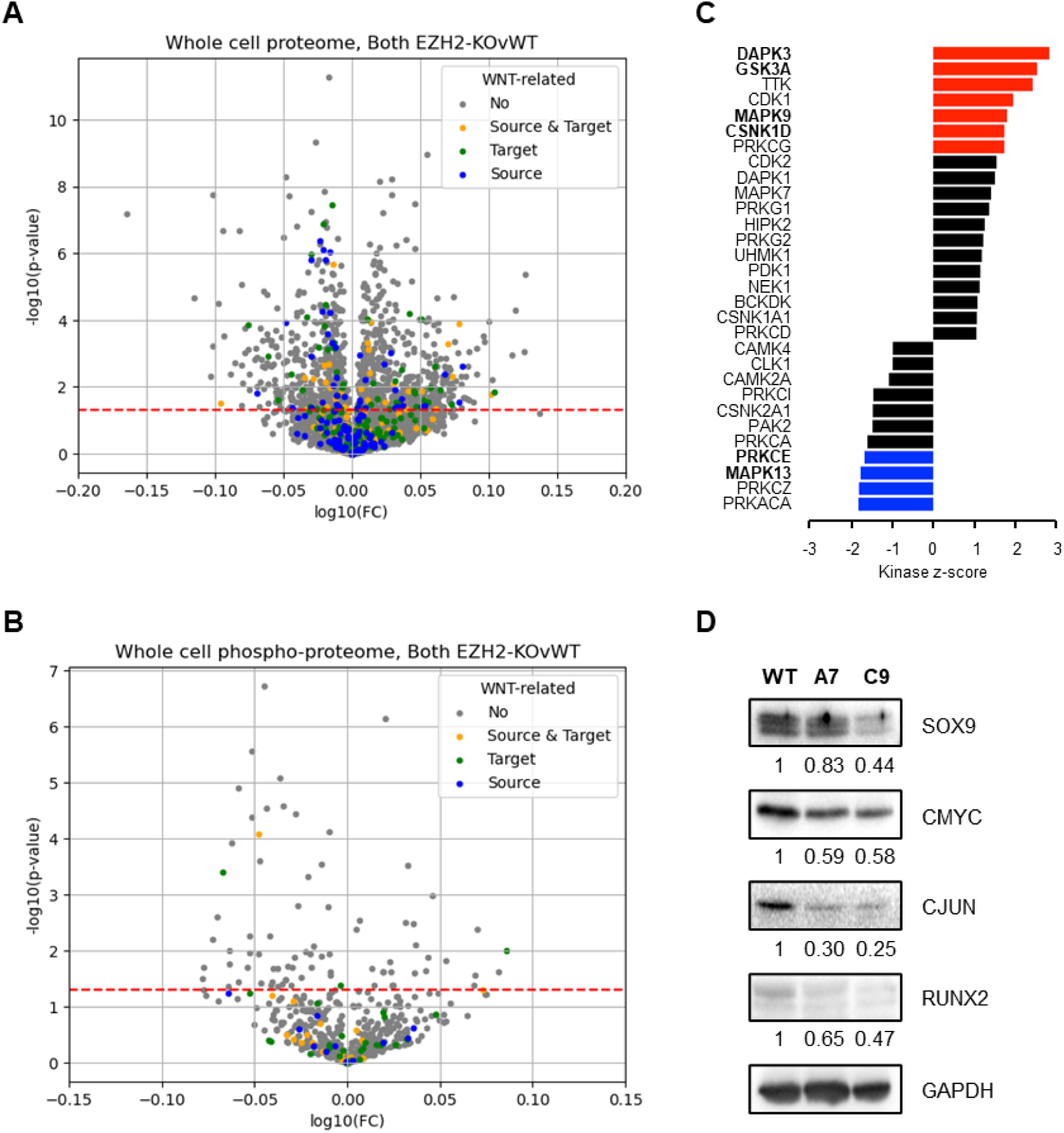
Multi-omics analysis reveals reduced WNT signaling activity in PRC2-depleted T-ALL. Volcano plots of **(A)** whole cell proteomic and **(B)** phosphoproteomic analysis of extracts from WT and EZH2-deficient cells. WNT pathways factors are highlighted in bold in each case. **(C)** Kinase Set Enrichment Analysis (KSEA, (22)) of phosphoproteomic outputs from WT and EZH2-KO cells. Kinases with relevance for WNT pathway activity are highlighted in bold. Colored bars indicate statistical significance. See Supplemental Figure S3B for comparison of individual EZH2-KO clones with WT samples. **(D)** Protein immunoblotting and quantification of WNT pathway targets.

Given the transcriptional results that suggested downregulation of WNT activity in isogenic EZH2-KO cells, we specifically examined proteomic changes in this pathway in our results. 398 WNT-related proteins were identified in our analyses, of which 61 were significantly differently expressed in either WT or EZH2-KO cells. Most of these (38/61) were down-regulated in both clones, with 18/61 commonly up-regulated, and only 5/61 proteins showed discordant changes between KO clones. WNT pathway factors that changed in expression between WT and KO cells are highlighted in Figure 3A and included PRKCA and RPS6KA1, which are known to modulate phosphorylation of negative WNT regulators GSK3α and GSK3β and hence downregulate WNT pathway signaling.

Phosphoproteomic profiling also revealed significant differences between WT and EZH2-null cells, including changes in WNT pathway-related factors, as highlighted in Figure 3B. To gain further insights into these results, we used the Kinase-substrate enrichment analysis (KSEA, (22)) tool that assigns a kinase activity score based on the phosphorylation status of known substrates using phospho-site-specific Kinase–Substrate (K–S) databases. Full details of these outputs are available in Supplemental Table S2 and are summarized in Figure 3C, with individual comparisons for each EZH2 clone shown in Supplemental Figure S3B. In line with our results so far, we found that significant KSEA changes included several kinases that are known to modulate WNT pathway activity. In particular, kinases that are known to inhibit WNT signaling (e.g., GSK3A, DAPK3, MAPK9, and CSNK1D) were found to have statistically significant increased activity by KSEA assessment, while kinases that have been described to stimulate WNT (e.g., PRKCE, MAPK13) had significantly lower KSEA scores.

Immunoblotting results also reflected decreased WNT activity in EZH2-depleted lines and were in line with proteomic and phosphoproteomic outputs, with protein levels of WNT transcriptional targets being markedly reduced in EZH2-null cells compared with their isogenic WT equivalents (Figure 3D).

### Asparaginase resistance in PRC2-deficient T-ALL is mediated by reduced WNT-STOP activity

Having established that PRC2 loss strongly correlates with reduced WNT pathway activity both *in vitro* and in patient samples, and in light of the previously reported links between PRC2 mutations and deletions and chemoresistance in acute leukemia (4–7), we next tested whether these signaling changes have any implications for T-ALL response to treatment.

We first assessed the responses of WT and PRC2-depleted Jurkat cells to a range of standard leukemia therapies, including all agents currently used in the remission induction phase of T-ALL treatment. Results were consistent with known drug sensitivities of the Jurkat cell line (23), with no observed differences in response to dexamethasone, daunorubicin, vincristine, or cytarabine between WT and EZH2-deleted cells (Supplemental Figure S4A). In contrast, we found a marked difference in sensitivity to asparaginase, with EZH2-null Jurkat cells being significantly more resistant to killing than their WT equivalents (p<0.001 in each case, Figure 4A and Supplemental Figure S4B).

**Figure 4:**
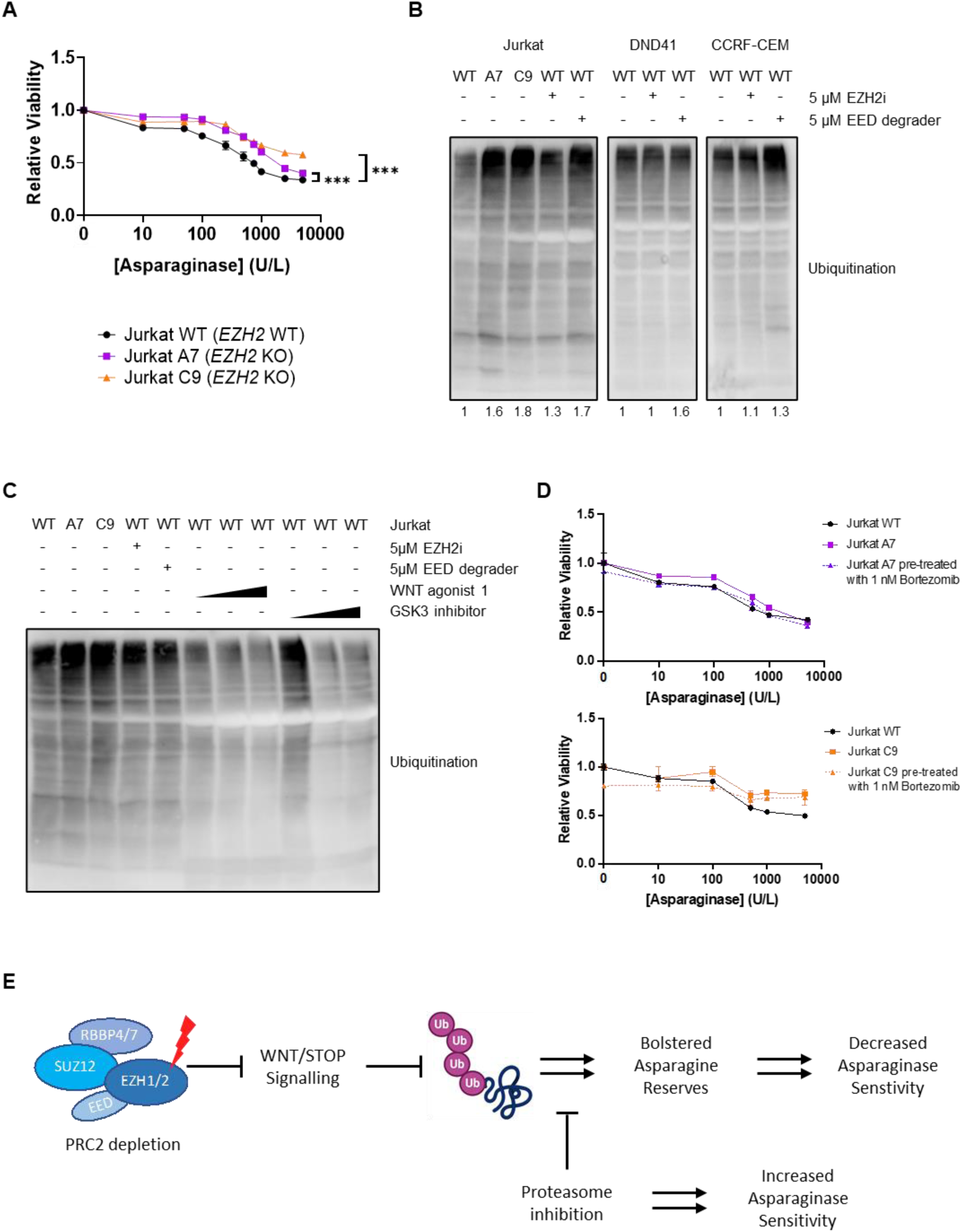
Asparaginase resistance in PRC2-deficient T-ALL is mediated by reduced WNT-STOP activity and can be mitigated by proteasome inhibition. **(A)** Differences in sensitivity of WT and EZH2-deficient Jurkat cells to asparaginase. A representative experiment is shown, with full results of all experimental replicates provided in Supplemental Figure S4B. Stars indicate statistical significance (p<0.001). **(B)** Measurement of ubiquitination levels by immunoblotting in (left to right) WT and EZH2-deficient Jurkat cells and other T-ALL cell lines (Jurkat, DND-41, and CCRF-CEM) treated for 48 hours with either enzymatic inhibition of EZH2 (GSK343) or a PROTAC EED degrader (UNC6852), as indicated. Quantification of ubiquitination relative to WT samples is shown under each blot. **(C)** Measurement of ubiquitination levels in T-ALL cell lines following treatment with either GSK343, UNC6852, a GSK3 inhibitor (BI-5521) or a WNT agonist molecule (WA1). **(D)** Comparison of single agent and combination treatment of proteasome inhibition (bortezomib) with asparaginase in WT and EZH2-KO Jurkat cell lines. A representative experiment is shown, with full results of all experimental replicates provided in Supplemental Figure S4G. **(E)** Proposed mechanism of asparaginase resistance through reduced WNT/STOP activity in PRC2-depleted T-ALL.

Recent reports have demonstrated a strong link between WNT pathway activity and asparaginase sensitivity in both acute leukemia and colorectal cancer (10,11) and led us to investigate whether this mechanism was responsible for asparaginase resistance in PRC2-depleted T-ALL. This described mechanism of resistance is caused by altered activity of the WNT/STOP pathway (24,25) that normally inhibits glycogen synthase kinase 3 (GSK3)-dependent protein ubiquitination and proteasomal degradation. Reduced WNT/STOP activity leads to increased protein ubiquitination and proteasomal degradation, thereby providing a catabolic source of amino acids that allows cancer cells to circumvent the asparagine depletion that is normally induced by asparaginase.

To explore this mechanism, we first measured protein ubiquitination in the Jurkat cell line model, and found markedly elevated levels of ubiquitination in EZH2-null cells (Figure 4B). We further confirmed that reducing PRC2 activity by either chemical EZH2 inhibition, or EED protein depletion using a PROTAC degrader, caused similar significant increases in protein ubiquitination in WT Jurkat and other T-ALL cell lines (Figures 4B and 4C, with confirmation of drug effect shown in Supplemental Figure S4C), whereas pharmaceutical stimulation of the WNT pathway with either a GSK3 stabilizer or WNT agonist caused decreased ubiquitination in T-ALL cells (Figure 4C). Of note, protein levels of asparagine synthetase (ASNS) were reduced in EZH2-null clones, suggesting that changes in ASNS activity did not contribute to asparaginase resistance in these cells (Supplemental Figure S4D).

Finally, we tested whether it was possible to modulate asparaginase resistance in PRC2-deficient cells using drug treatments. We first tried to stimulate WNT pathway signaling using either a GSK3 stabilizer or WNT agonist, but found that these agents caused significant toxicity in T-ALL cell lines that precluded assessing their effectiveness in combination with asparaginase (Supplemental Figure S4E).

We then hypothesized that treatment with a proteasome inhibitor would counteract the asparaginase resistance observed in PRC2-deficient cells, by depleting the catabolic source of asparagine provided by proteasomal protein degradation. As proteasome inhibition is toxic to T-ALL cells, we used a very low dose of the proteasome inhibitor bortezomib (1nM) that did not cause cell death when administered as a single agent (Supplemental Figure S4F) in our combination experiments. In line with our hypothesis, we found that bortezomib treatment mitigated asparaginase resistance in EZH2-null Jurkat cells, in each case at least partially restoring sensitivity to the levels observed in the WT line (Figure 4D and Supplemental Figure S4G), thereby confirming a direct role for proteasomal activity in this resistance mechanism, and providing a potential therapeutic avenue to increase asparaginase sensitivity *in vivo*.

## Discussion

In this study, we provide a holistic analysis of signaling pathway changes in PRC2-depleted T-ALL, identify a mechanistic explanation of resistance to a key blood cancer treatment, and suggest potential solutions to overcome this important clinical problem.

Asparaginase is a central plank of leukemia treatment during the initial phase of remission induction therapy, and suboptimal asparaginase responses because of intrinsic cellular resistance or acquired immune sensitivities are strongly linked to poor outcomes. Why some T-ALLs fail to respond to this agent is not fully clear. Sensitivity is at least partly mediated by levels of asparagine synthetase (*ASNS*) expression and epigenetic modification of *ASNS* transcription, but these relationships are inconsistent. In line with this, EZH2-depleted cells in our experiments had decreased levels of ASNS that would be predicted to cause increased asparaginase sensitivity, but we found that the opposite was the case.

Recent insights into asparaginase resistance mechanisms were provided by several elegant reports that linked WNT/STOP pathway activity to asparaginase sensitivity in leukemia and colorectal cancer (10,11). Our results add to this knowledge by providing a novel link between asparaginase resistance by this mechanism and a genetic alteration in T-ALL cells, i.e., reduced Polycomb function through mutation or deletion of core PRC2 factors. To our knowledge, this is the first description of a mechanistic link between a somatic genetic alteration and response to a specific treatment agent in T-ALL.

These investigations were triggered by our identification in this study of previously unreported genetic associations between PRC2 factors and signaling factor molecules, thereby highlighting the value of careful examination of mutual exclusivities and co-occurrences within genomic data. While previous studies in this area primarily focused on mutations, our analyses included both mutations and copy number alterations, thereby allowing us to identify these novel associations.

One limitation of our study was the cellular context in which much of the molecular analysis of signaling pathway activity was carried out. Whereas most PRC2 mutations and deletions in patient cases occur in the immature/ETP-ALL subgroup, the Jurkat cell line corresponds to later differentiation arrest at a cortical stage of T-lymphoid development. Our attempted deletion of *EZH2* in other T-ALL cell lines was unsuccessful (data not shown), which might reflect gene essentialities in these models.

Despite this caveat, it is important to note that previous analyses showed increased expression of genes associated with an immature/ETP-ALL phenotype in this model (20), and which therefore would mimic the most frequent transcriptional setting in patient cells. In addition, identification of a specific molecular vulnerability to CHK1 inhibition in this experimental system was corroborated by results in PRC2-altered primary patient samples, providing support for this model as a reliable proxy for PRC2-altered T-ALL *in vivo*. Further, our key finding that transcription of WNT pathway genes is reduced in PRC2-altered patient leukemia samples (Figure 2F), in line with expression patterns seen in our *in vitro* model (Figure 2F), and the fact that drug targeting of PRC2 markedly increases ubiquitination activity in other cell lines (Figure 4B), supports our confidence that these findings can be extrapolated to other T-ALL contexts.

We have shown that co-administration of the proteasome inhibitor bortezomib can mitigate asparaginase resistance in PRC2-depleted cells, thereby suggesting a potential avenue towards modulating therapy response in patients in future. We would like to highlight that the choice of dose used in our experiments was based on careful titration of bortezomib concentration to non-toxic levels (Supplementary Figure S4D), in order to avoid any confounding effects of direct bortezomib activity on cell viability results. In our experiments, even this very small dose of 1nM resulted in improved asparaginase sensitivity in combination treatments. It is highly likely that uses of even moderately higher doses of proteasome inhibitor would result in increased levels of leukemia cell death that could be of greater benefit in the clinical setting.

A key question raised by our findings is how PRC2 and other epigenetic factors might regulate the expression of genes that modulate the activity of WNT and other signaling pathways in leukemia cells. While our results provide a profile of the transcriptional changes linked to EZH2 loss, an important next step will be to analyze how PRC2 mutations and deletions affect the binding of Polycomb and other epigenetic factors across the genome, and to WNT-related loci in particular. It will also be critical to evaluate whether other T-ALL somatic genetic changes also alter WNT/STOP activity, and whether co-administration of proteasome inhibitors might provide opportunities to improve asparaginase responses and overall treatment outcomes in other high-risk leukemia subgroups.

## Acknowledgements

The authors thank the National Children’s Research Centre for their significant support of this work through Leadership Award A-18-3 that funded start-up research in the Bond laboratory at Systems Biology Ireland, and which fully supported TL’s doctoral work. The authors would also like to thank and acknowledge the many team members at Systems Biology Ireland and the Irish National Children’s Cancer Service at Children’s Health Ireland at Crumlin for their support and helpful suggestions during the course of this project.

Work in the Bond laboratory is also funded by Science Foundation Ireland grants 20/FFP-P/8844 and 18/SPP/3522, the latter together with Children’s Health Ireland. Work in the Ryan laboratory is supported by Science Foundation Ireland grant 20/FFP-P/8641. CT was funded by Science Foundation Ireland through the SFI Centre for Research Training in Genomics Data Science under Grant number 18/CRT/6214 and supported in part by the EU’s Horizon 2020 research and innovation programme under the Marie Skłodowska-Curie grant H2020-MSCA-COFUND-2019-945385. Proteomic and phosphoproteomic analyses were supported by the Comprehensive Molecular Analytical Platform (CMAP) under the SFI Research Infrastructure Programme, reference 18/RI/5702.

## Methods

### Mutational co-occurrence analysis

Mutational and copy number alteration data from a cohort of childhood, adolescent, and young adult patients with T-ALL (2) were analyzed using the Discrete Independence Statistic Controlling for Observations with Varying Event Rates (DISCOVER) pipeline (13). Full details are provided in the supplemental methods.

### CRISPR/Cas9 gene editing

2 guide RNAs (gRNAs, gRNA#1: CGGAAATCTTAAACCAAGAATG, gRNA#2: ACCAAGAATGGAAACAGCGAAG) were designed to direct the Cas9 endonuclease to exon 3 of *EZH2*. gRNAs were purchased from Integrated DNA Technologies as part of the Alt-R CRISPR-Cas9 System that delivers the full ribonucleoprotein (RNP) complex of gRNA, tracrRNA and Cas9 enzyme. Jurkat cells were transfected using the Cell Line Nucleofector™ Kit V (Lonza, VCA-1003) coupled with the Amaxa® Nucleofector® System (Lonza, Nucleofector 2b).

### RNA-sequencing

RNA was extracted using the RNAeasy kit (Qiagen). PolyA-enriched paired-end sequencing was performed using the Illumina NovaSeq 6000 Sequencing System at a depth of 30 million reads. Workflows for pre-processing and analyzing RNA-sequencing data to generate gene counts were derived from https://diytranscriptomics.com/ and is available on request. PROGENy (Pathway RespOnsive GENes) was performed according to the original description of the pipeline (21) available at https://github.com/saezlab/transcriptutorial. GSEA using the H, C2 and C6 sets was performed using the GSEABase 1.56.0 and msigdbr 7.4.1 R packages.

### Proteomic analysis

Mass spectrometry (MS) output data was searched using MaxQuant (release 2.0.1.0) against the *Homo sapiens* subset of the Uniprot Swissprot database with specific parameters for TIMS data dependent acquisition (TIMS-DDA). The MaxQuant output file was imported into the Perseus (version 1.6.15.0) environment for protein quantification.

Phosphoproteome data was similarly exported and transformed to enable analysis of phosphorylation level and residue position. Exported data were loaded into R environment (R version 4.0.4, libraries: dplyr 1.0.8, gtools 3.9.2) for further analysis. Kinase-Substrate Enrichment Analysis (KSEA) was performed as previously described (22).

Full methods for cell culture, assessment of *EZH2-*deleted Jurkat clones, mass spectrometry, immunoblotting, drug treatments, cell viability testing, RNA-seq data analysis, and KSEA are provided in the **Supplemental Methods**.

## Supplemental Figure S1 (Corresponds to Figure 1): **Results of DISCOVER analysis.**

**Figure S1A:**
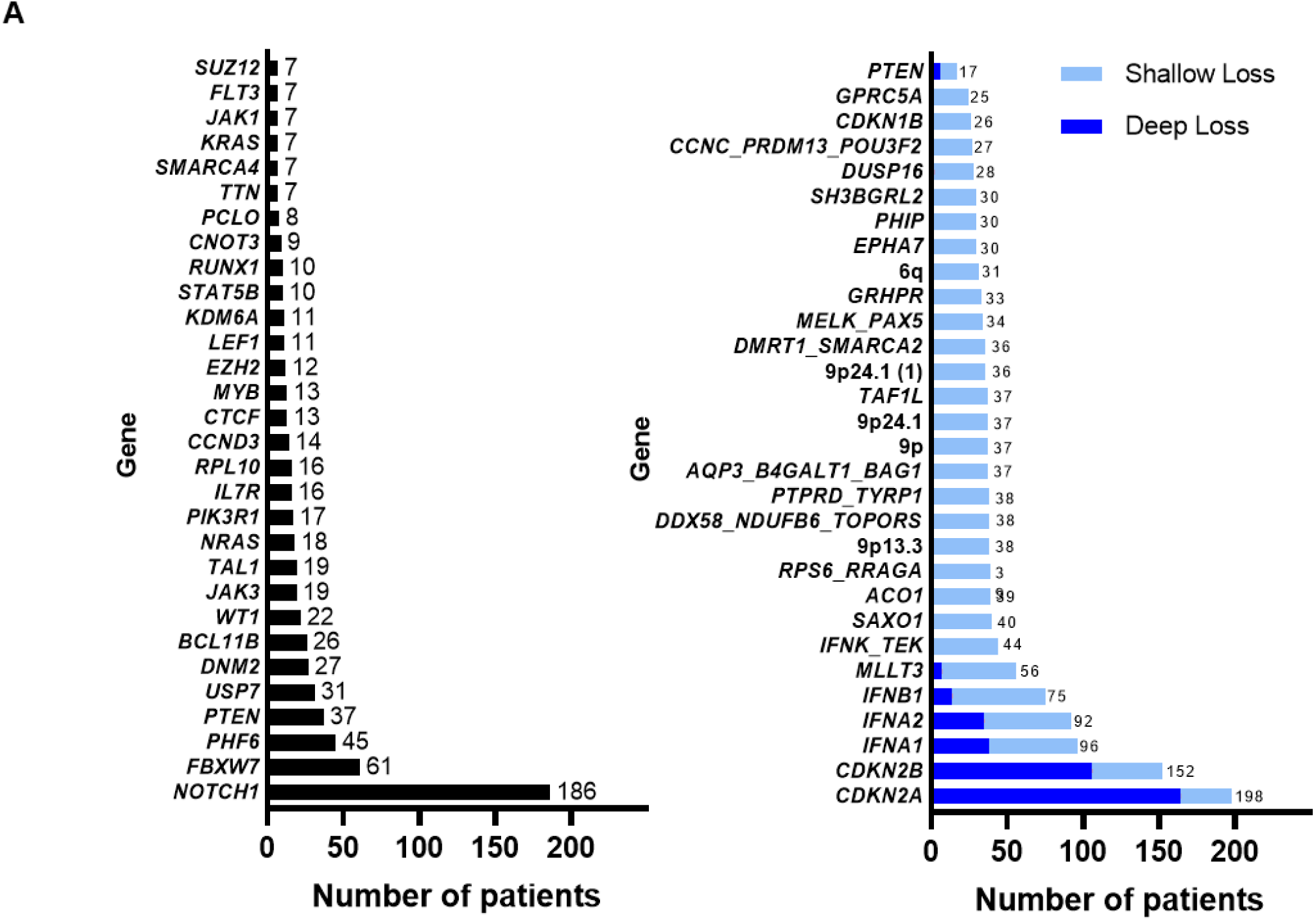
Top 30 genes with highest frequency of mutations (left) and top 30 loci with copy number alterations (right) in the pediatric T-ALL cohort (13) used for DISCOVER analysis.

**Figure S1B:**
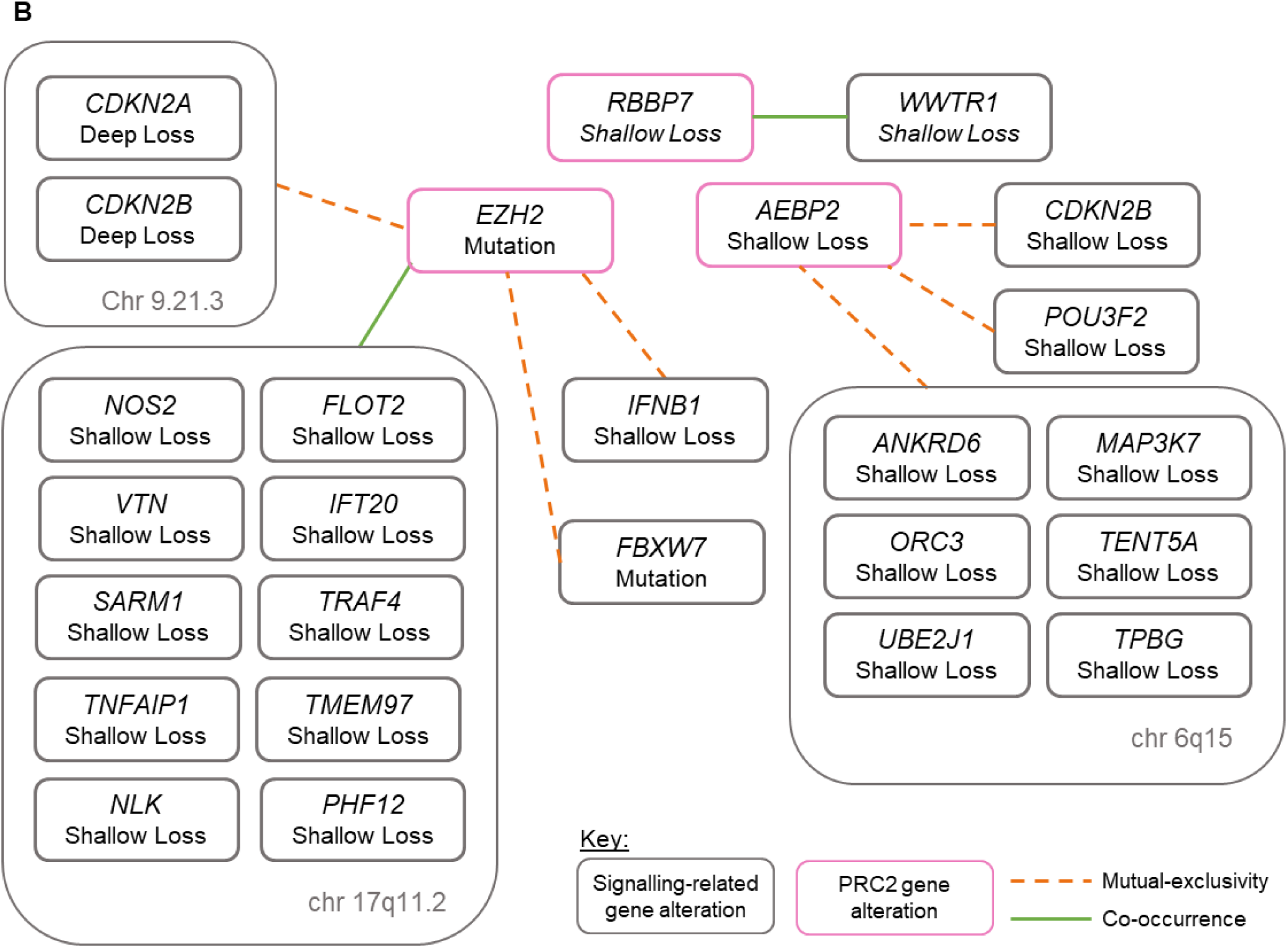
Graphical representation of statistically significant interactions between genes coding for epigenetic factors (pink boxes) and signaling pathway components (grey boxes) in the DISCOVER analysis. Mutual exclusivity and co-occurrence relationships are indicated.

## Supplemental Figure S2 (Corresponds to Figure 2): **Loss of PRC2 function correlates with reduced transcription of WNT pathway targets.**

**Figure S2A:**
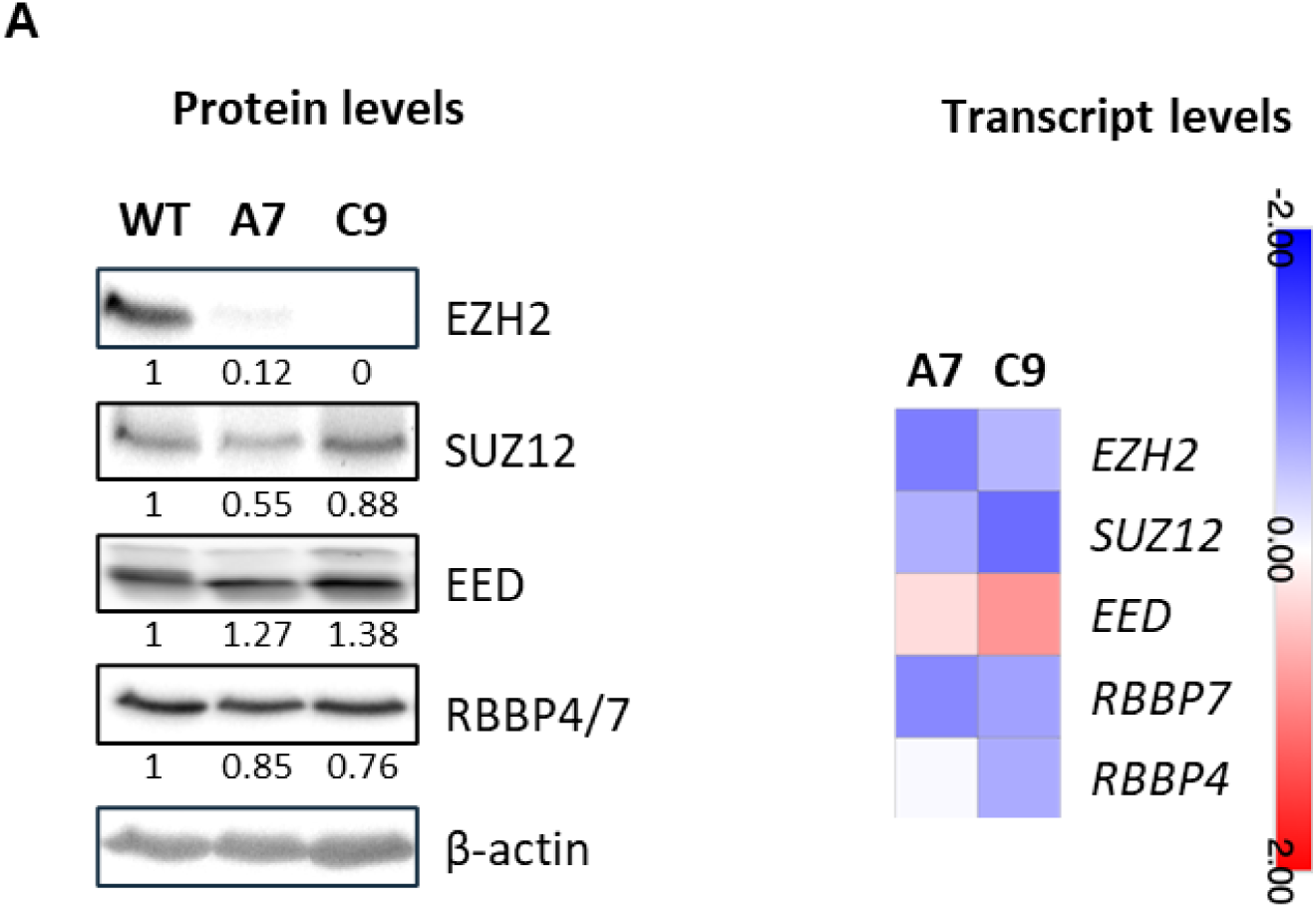
Measurement of protein levels of PRC2 components by immunoblotting with quantification results under each image (left panel), with quantification of the corresponding mRNA transcript by RNA-sequencing in the accompanying heatmap (right panel). In each case, expression levels in the EZH2 KO clones were normalized to the WT samples.

**Figure S2B:**
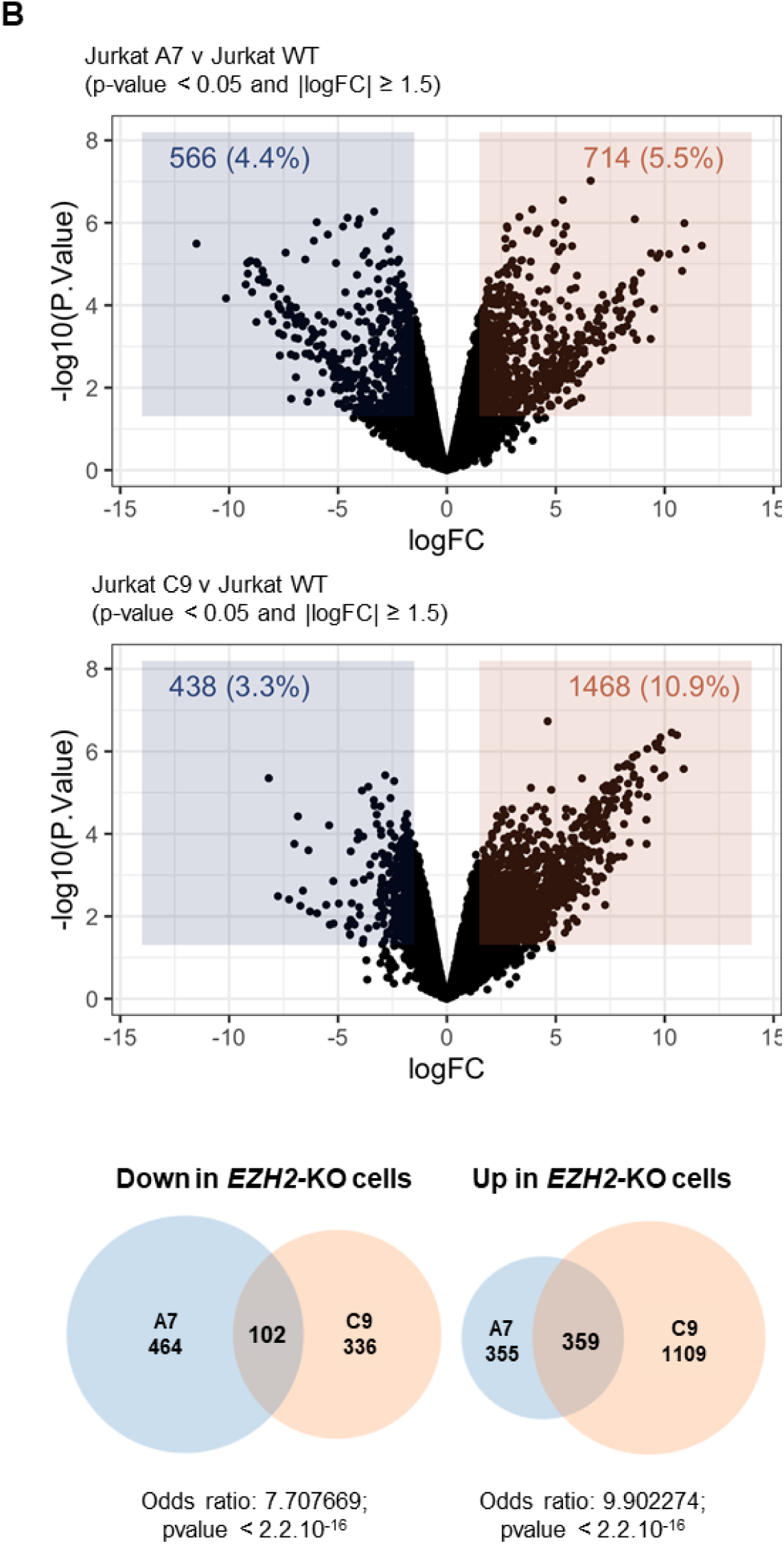
Volcano plots depicting differential gene expression in EZH2-null clones A7 (top panel) and C9 (middle panel) compared with WT. The bottom panel shows a Venn diagram of the observed transcriptional differences between each PRC2-depleted clone and WT cells, and statistical analysis of the overlap in measured gene expression changes using Fisher’s exact test.

**Figure S2C:**
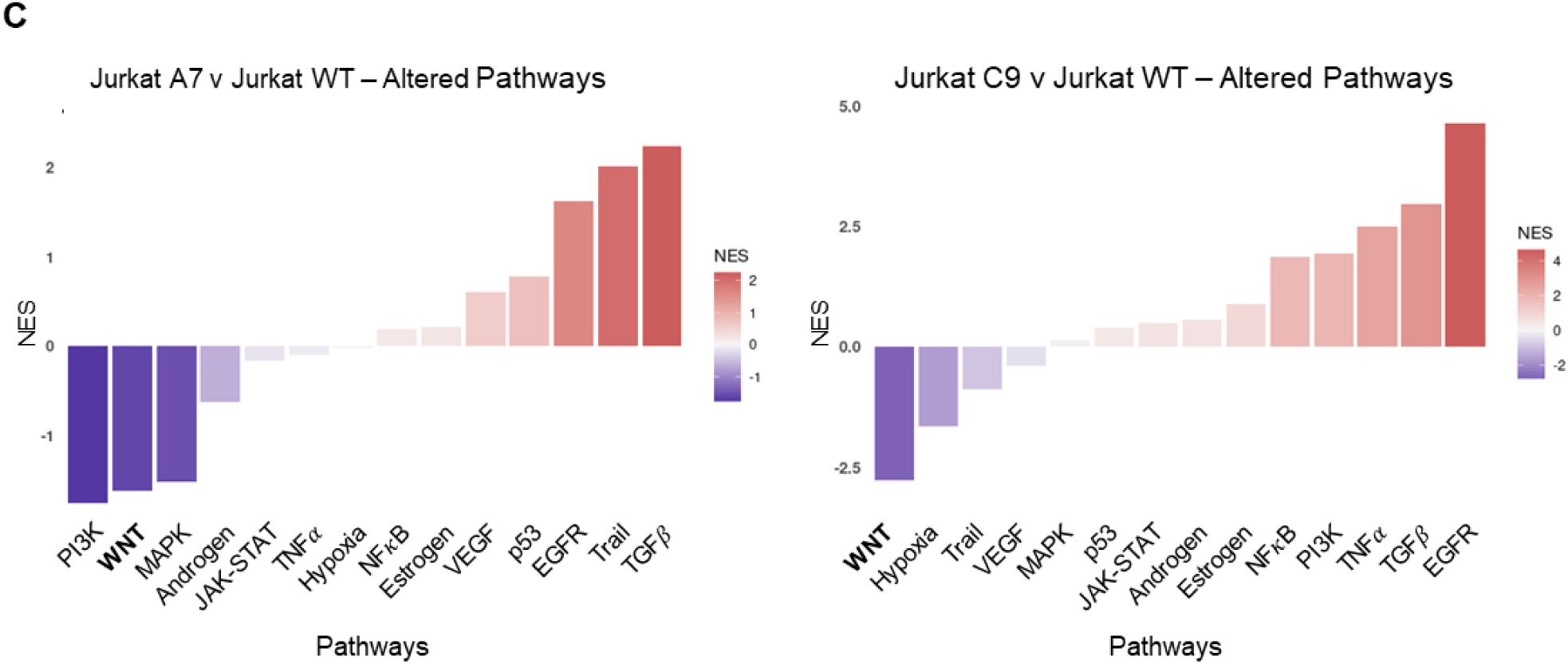
PROGENy analysis of signaling pathway activity from RNA-seq transcriptional readouts in individual clones (see Figure 2C for combined analysis of clones v WT samples). Enrichment in EZH2-KO cells is depicted, i.e., negative enrichment corresponds to predicted decreased activity of that pathway in PRC2-depleted cells. NES: Normalized enrichment score.

**Figure S2D:**
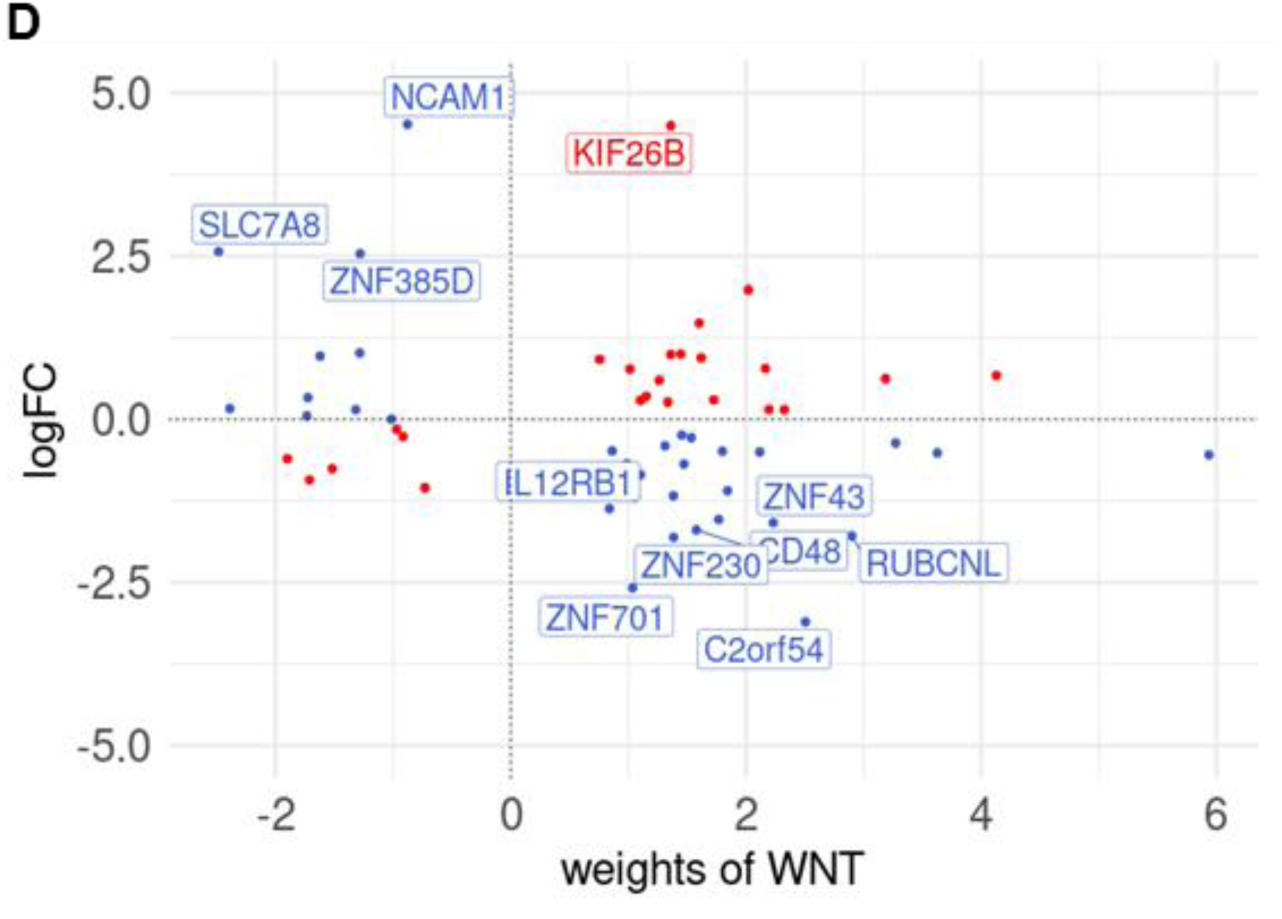
Full results for transcriptional changes in PROGENy WNT pathway targets. Log Fold Change (logFC) in expression level is shown on the y axis, with scoring weights of each factor on the x axis. Genes shown in the heatmap in Figure 2D in the main results section are highlighted.

**Figure S2E:**
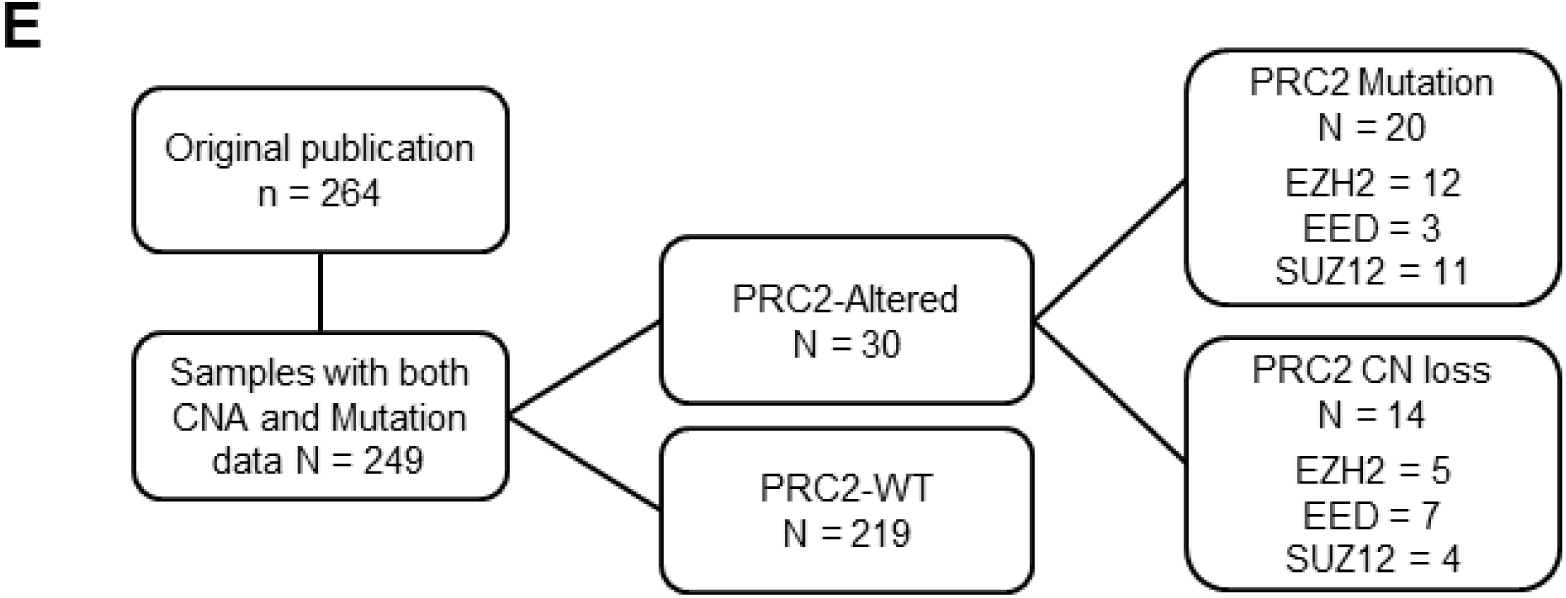
Outline of the pediatric T-ALL cohort (13) analyzed in Figure 2F. Numbers of cases with mutations and deletions (CN = copy number) in core PRC2 factors are shown. 4 cases had co-occurrence of mutation and deletion in PRC2 genes.

**Figure S2F:**
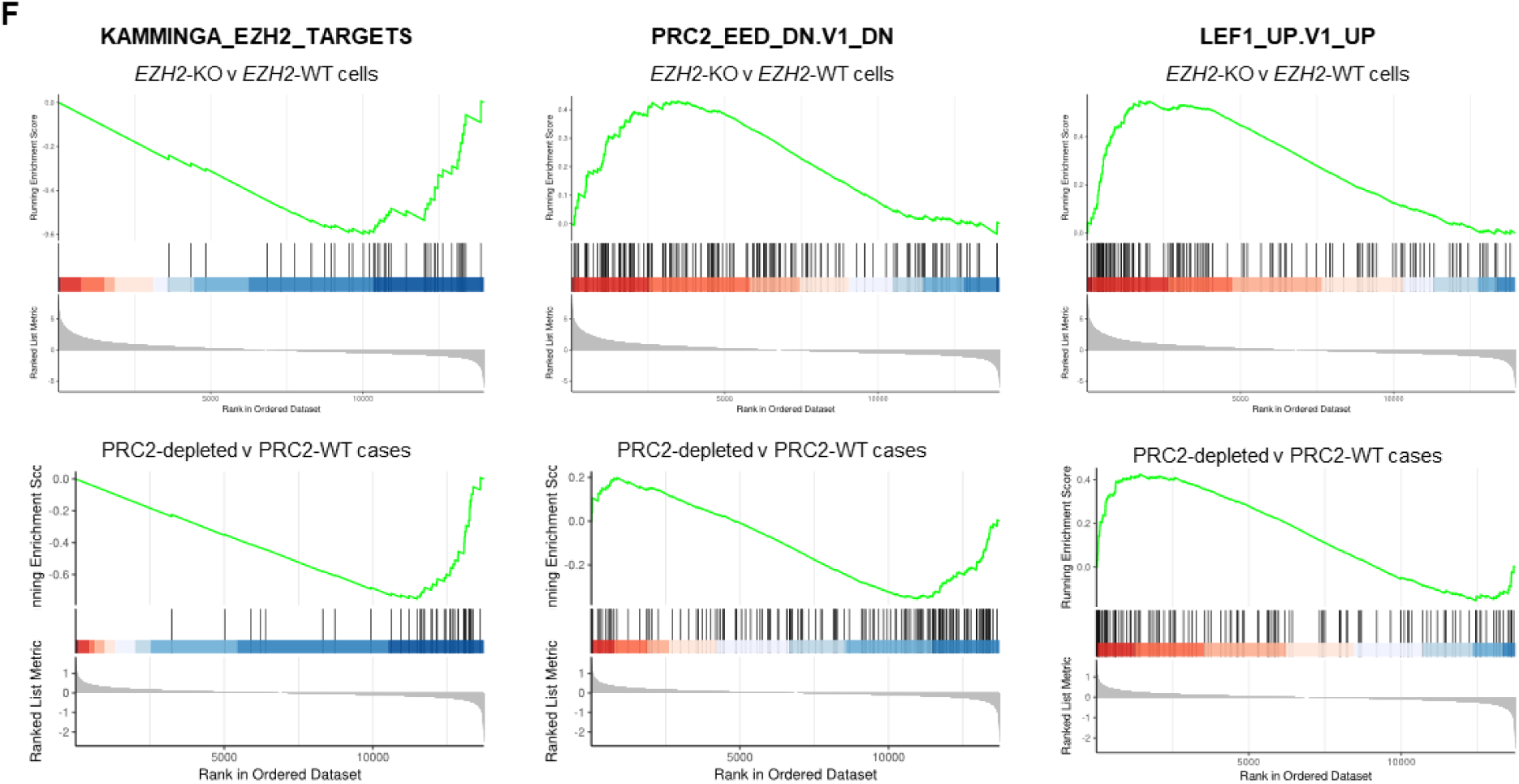
Gene Set Enrichment Analyses (GSEA) of the Jurkat cell line model (top 3 panels) and the T-ALL patient cohort (bottom 3 panels). Gene sets used for GSEA are indicated above the plots and correspond to previously described targets of PRC2 components EZH2 and EED, and WNT effector LEF1. In each case, transcriptional enrichment in PRC2-depleted Jurkat cells is consistent with the patterns observed in PRC2-altered primary T-ALL samples.

## Supplemental Figure S3 (Corresponds to Figure 3): **Comparison of protein expression changes between Jurkat WT and EZH2-null clones.**

**Figure S3A:**
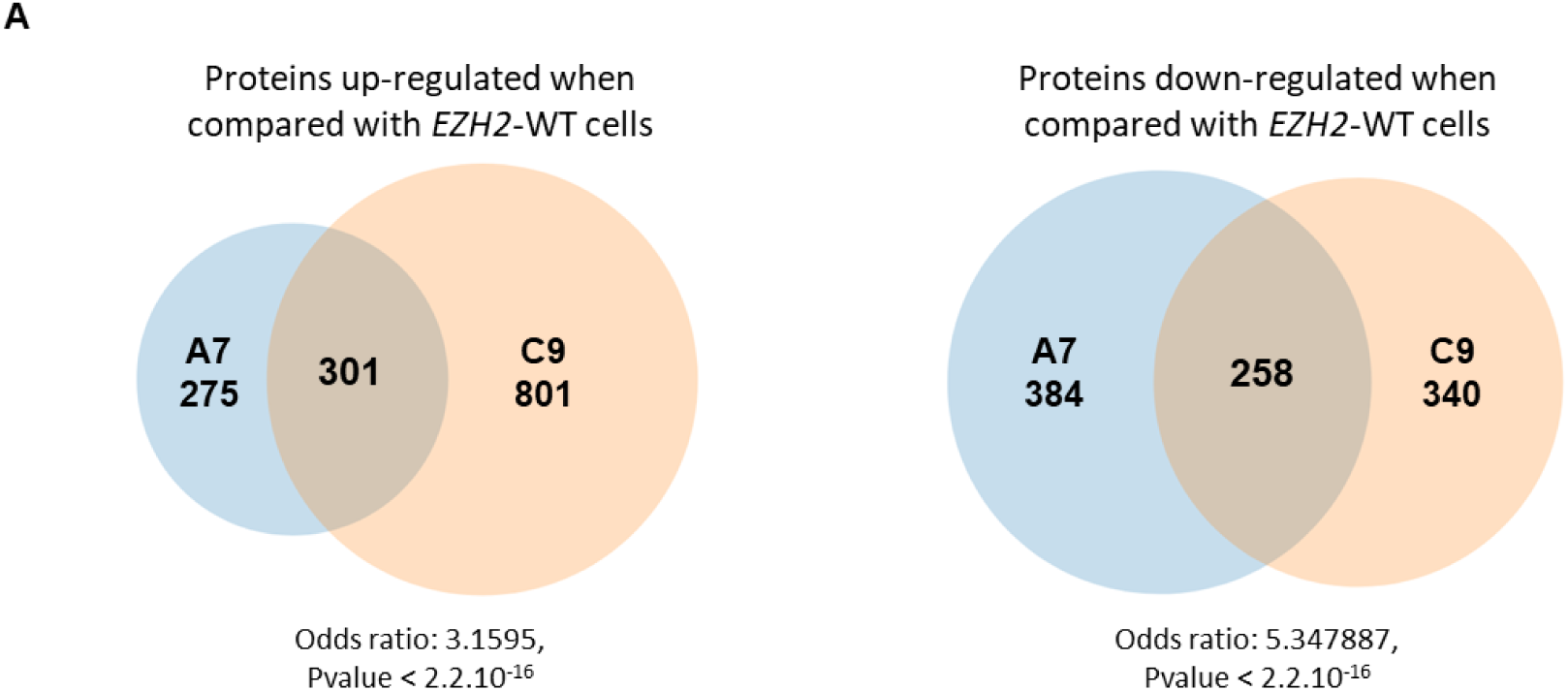
Venn diagram showing overlap of all measured upregulated and downregulated proteins in EZH2-null Jurkat clones A7 and C9 compared with WT, as determined by mass spectrometric assessment. Results of statistical analysis of the overlap in protein level changes by Fisher’s exact test are shown.

**Figure S3B:**
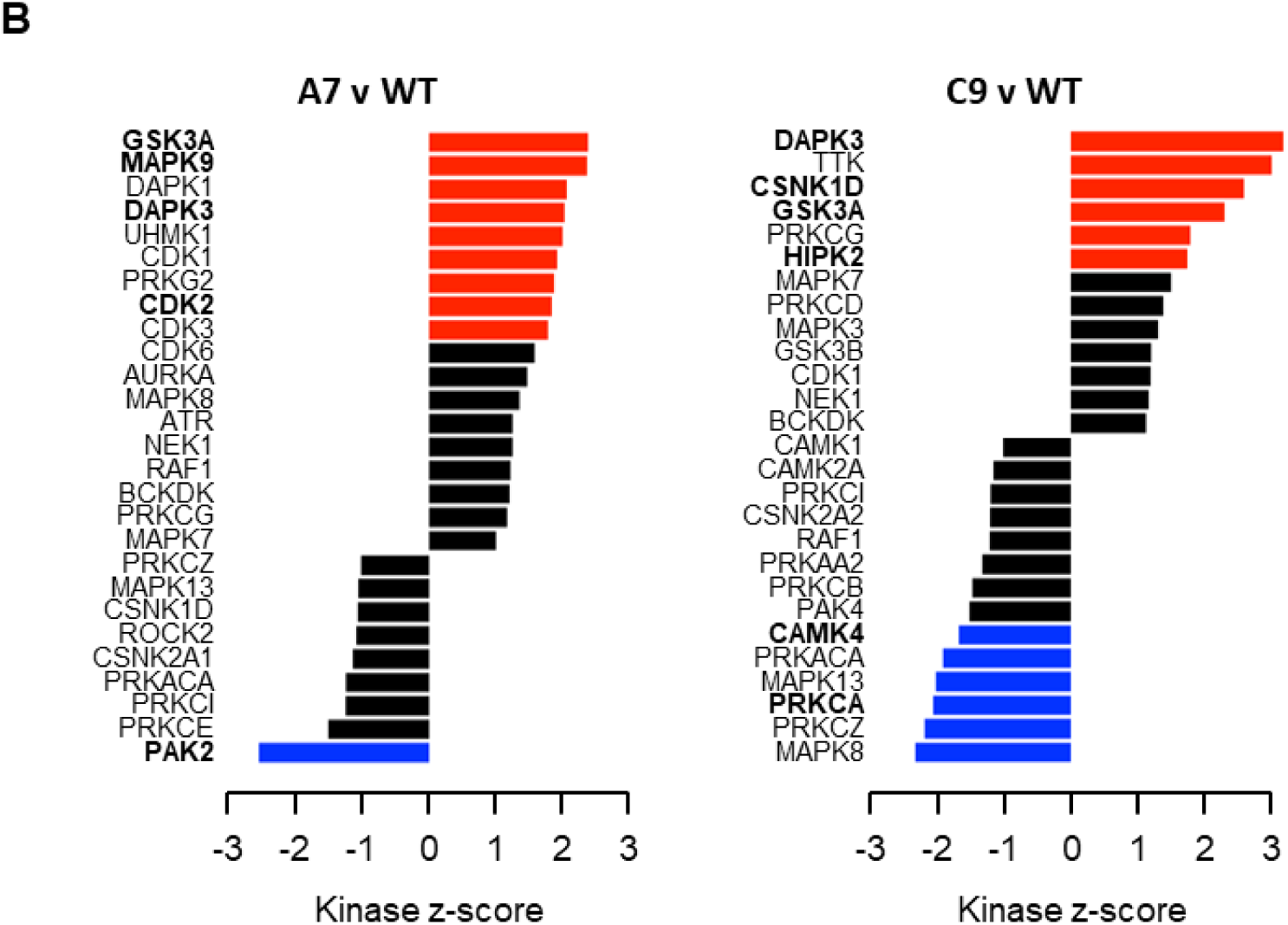
Kinase Set Enrichment Analysis (KSEA) of individual clones compared with WT samples. Kinases with relevance for WNT pathway activity are highlighted in bold. Colored bars indicate statistical significance. See Figure 3C for combined analysis of clones v WT samples.

## Supplemental Figure S4 (Corresponds to Figure 4): **Asparaginase resistance in PRC2-deficient T-ALL is mediated by reduced WNT-STOP activity and can be mitigated by proteasome inhibition.**

**Figure S4A:**
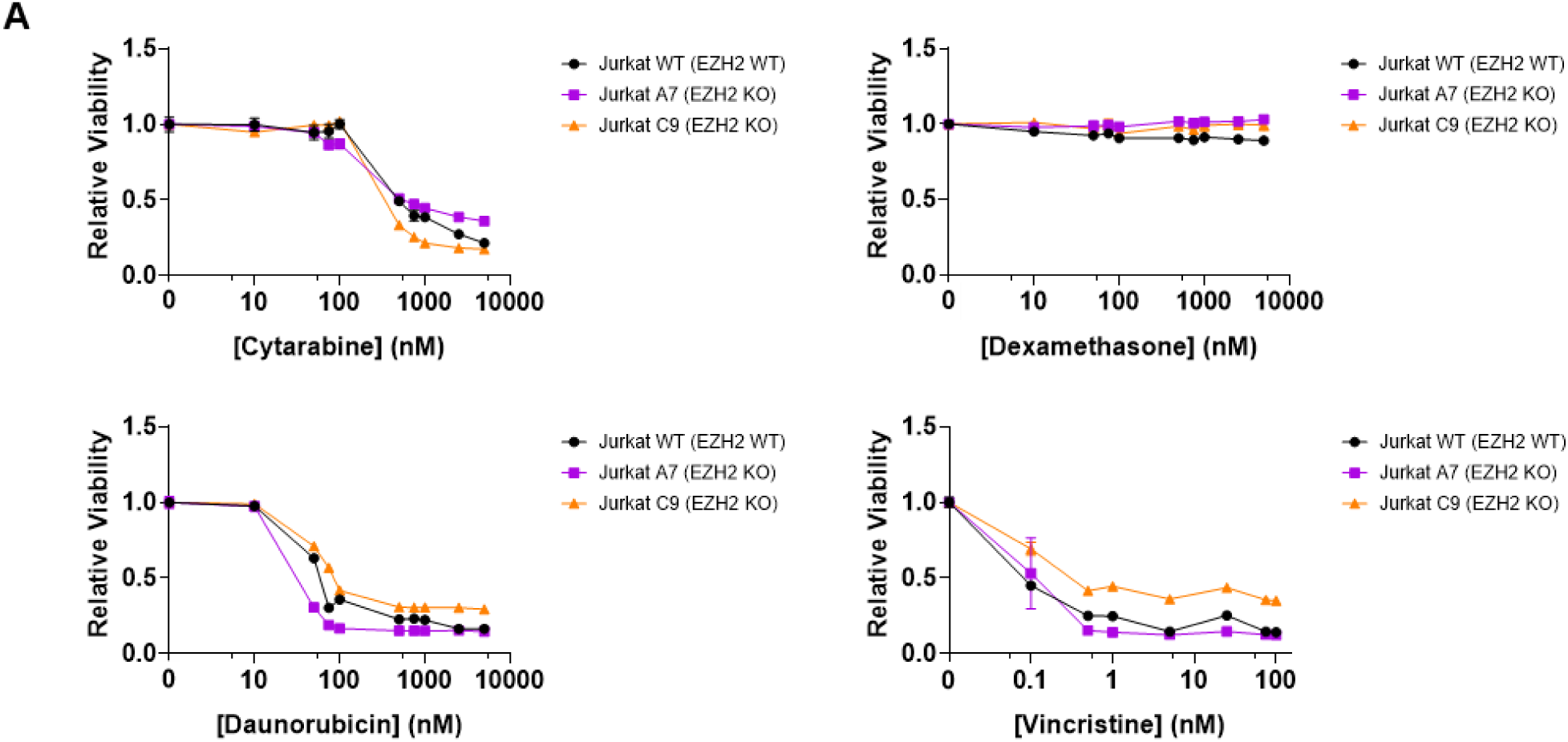
Comparison of sensitivities of WT and EZH2-deficient Jurkat cells to dexamethasone, daunorubicin, vincristine, and cytarabine. Results are consistent with reported patterns of response in WT cells, including corticosteroid resistance (14).

**Figure S4B:**
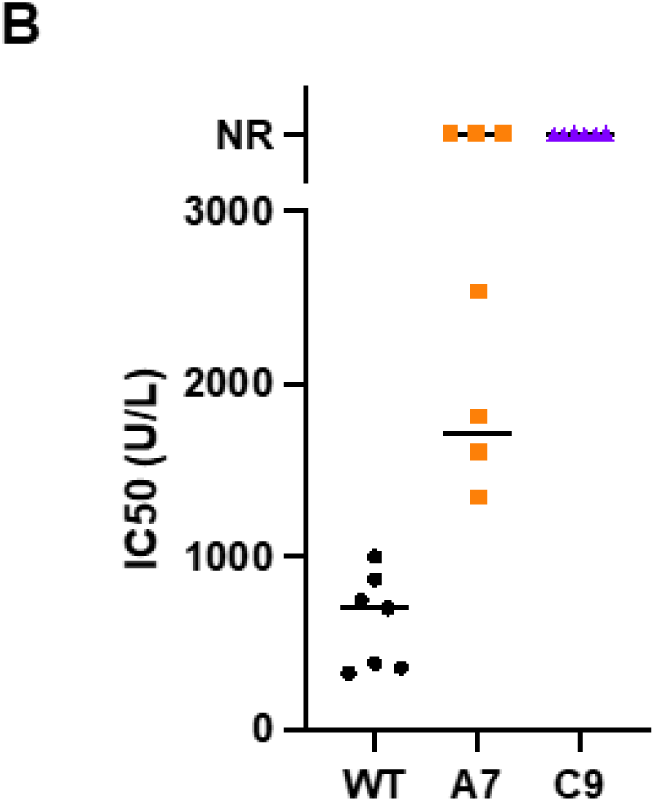
Full results of experimental replicates of asparaginase treatment of WT and EZH2 KO clones. Asparaginase doses required to reach 50% cell death (IC50) are shown. NR = not reached, i.e., 50% cell death was not attained in that experiment.

**Figure S4C:**
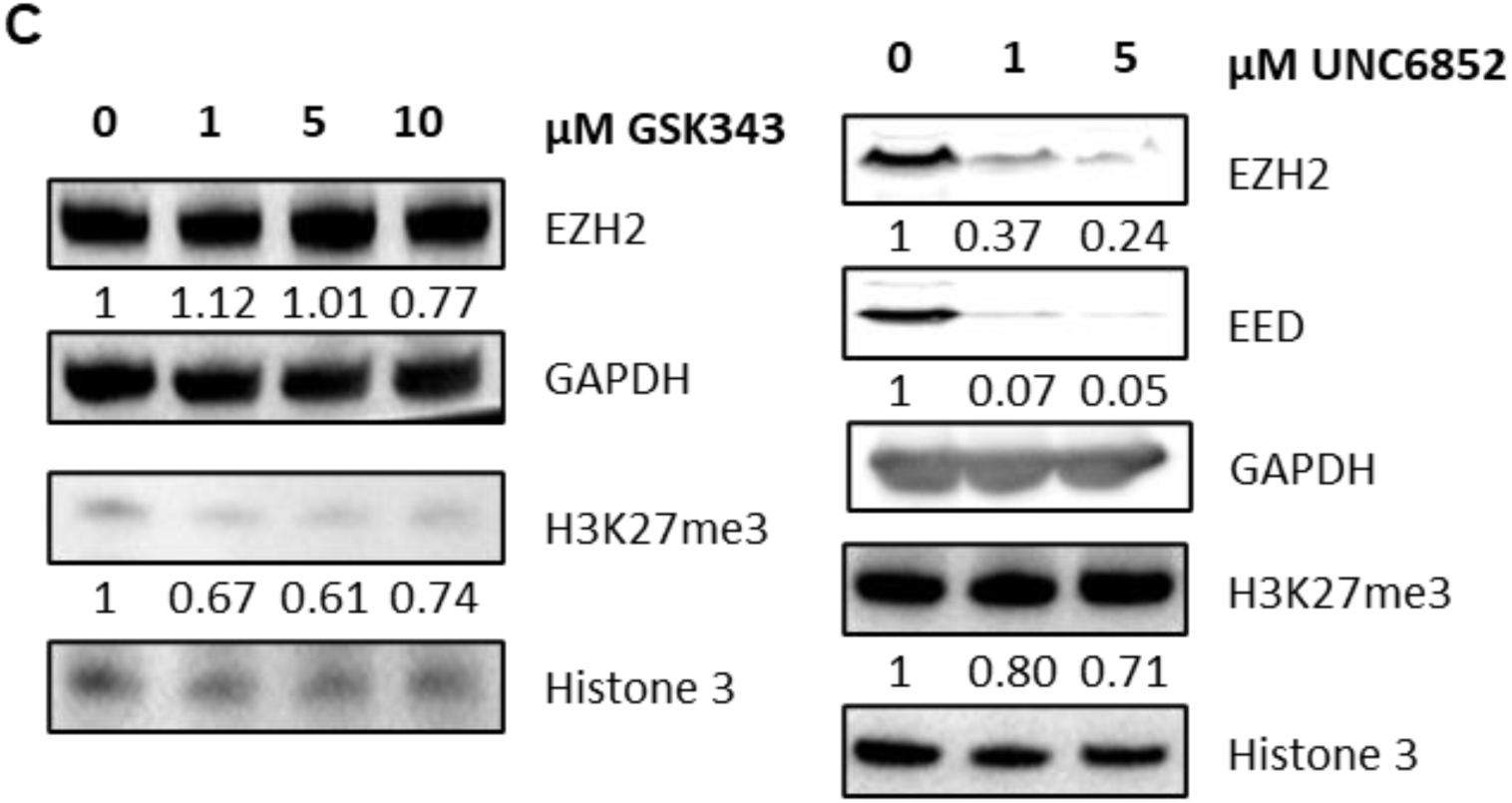
Effects of treatment of Jurkat T-ALL cells with an enzymatic EZH2 inhibitor (left panel) and PRC2 degrader (right panel). Immunoblots show reductions in H3K27me3 in each case, and depletion of EZH2 and EED proteins in the case of the UNC6852 treatment. Quantification relative to a loading control is shown below the relevant blots.

**Figure S4D:**
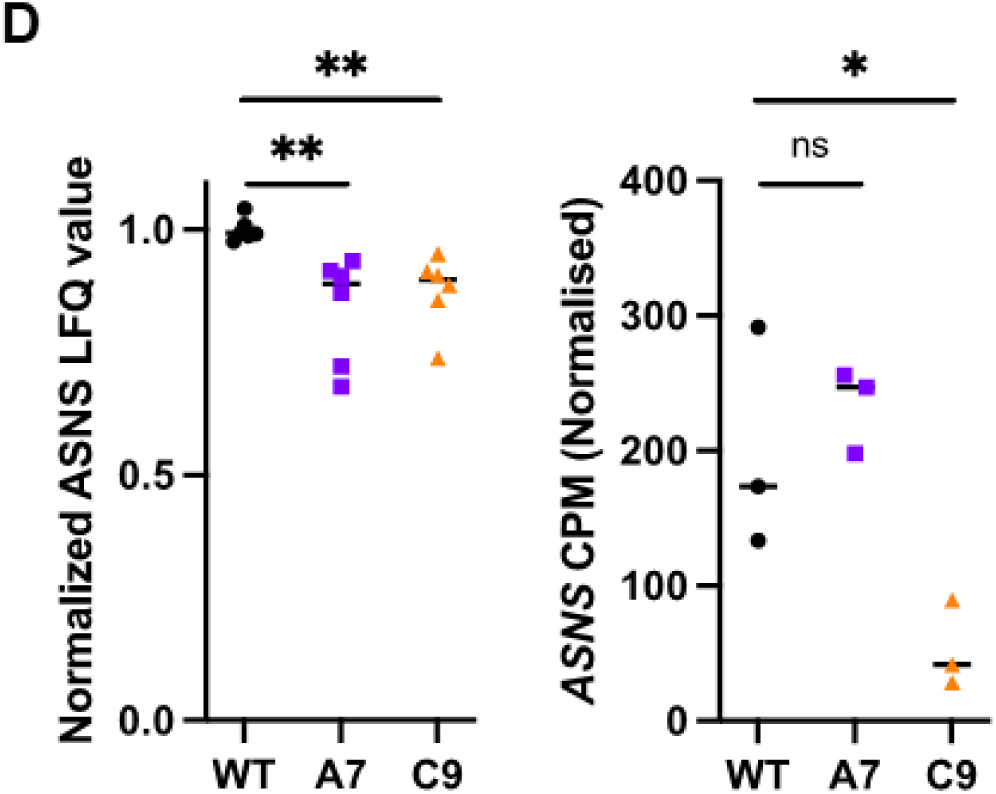
Comparison of asparagine synthetase (ASNS) protein and mRNA levels in WT and EZH2 KO cells. Left panel: protein levels as measured by mass spectrometry. Right panel: mRNA levels as measured by RNA-sequencing. Statistically significant differences are indicated.

**Figure S4E:**
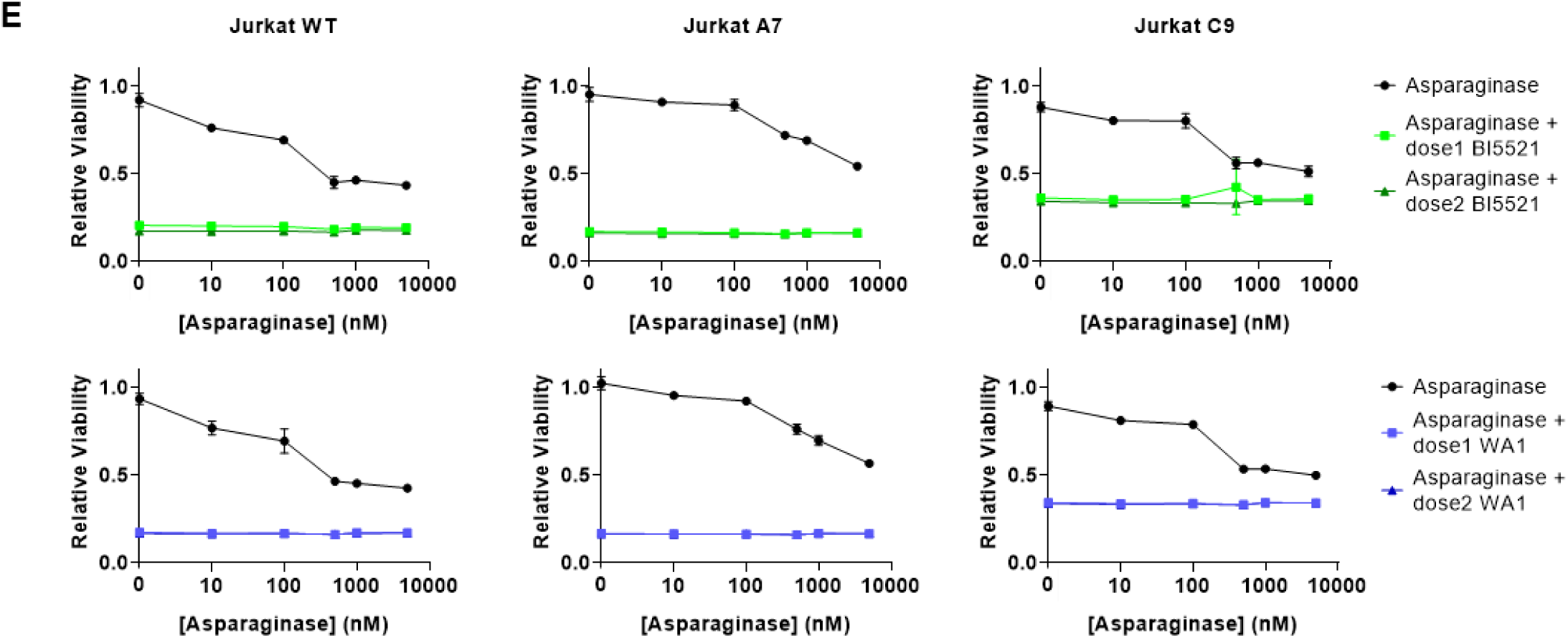
Effects of co-administration of asparaginase and a GSK3 stabilizer (top panels) or a WNT agonist (bottom panels) showing significant toxicity of each drug that precluded assessing the effectiveness of this combination strategy.

**Figure S4F:**
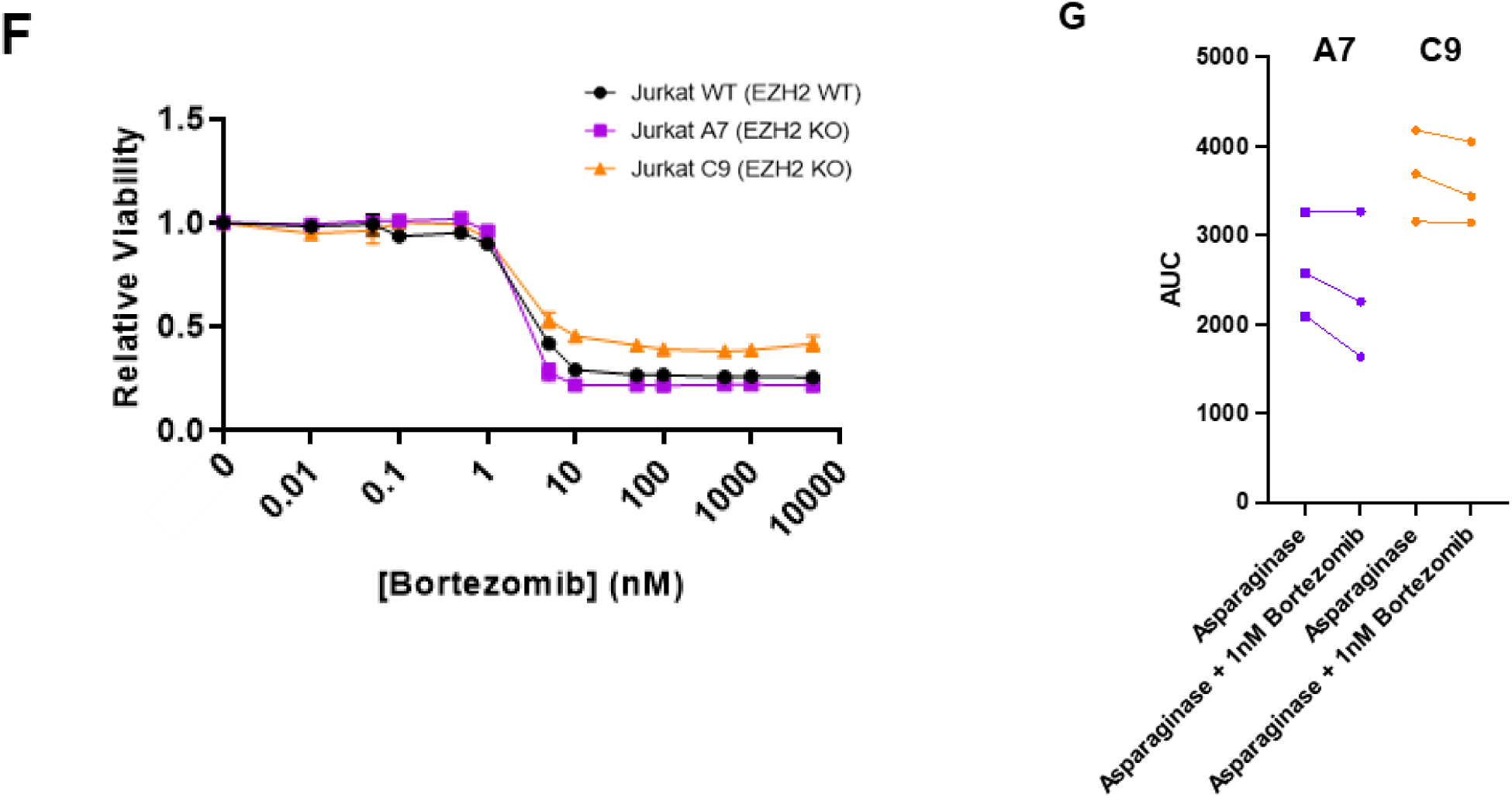
Titration of proteasome inhibitor bortezomib treatment to determine the threshold of non-toxic dosage levels. A concentration of 1nM (i.e., the highest dose that was not toxic to Jurkat T-ALL cells in single agent treatment) was used in subsequent combination experiments with asparaginase.

**Figure S4G:** Individual replicates for combination treatments of asparaginase with the non-toxic dose of 1nM bortezomib. Treatments carried out in the same experiment are linked by bars. AUC = Area under the curve.

